# Mitochondrial interactome quantitation reveals structural changes in metabolic machinery in failing murine heart

**DOI:** 10.1101/2021.08.13.456027

**Authors:** Arianne Caudal, Xiaoting Tang, Juan D. Chavez, Andrew Keller, Outi Villet, Bo Zhou, Matthew A. Walker, Rong Tian, James E. Bruce

**Author notes:** Co-first author (Arianne Caudal, Xiaoting Tang). Correspondence to: James E. Bruce, PhD, Department of Genome Sciences, University of Washington, 850 Republican Street, Seattle, WA 98109, Rong Tian, MD, PhD, Mitochondria and Metabolism Center, University of Washington, 850 Republican Street, Seattle, WA 98109.

## Abstract

Advancements of cross-linking mass spectrometry (XL-MS) for structural analysis of proteins bridges the gap between purified systems and native tissue environments. Here, isobaric quantitative protein interaction reporter technology (iqPIR) was utilized to further extend XL-MS to the first system-wide comparative study of mitochondrial proteins from healthy and diseased murine hearts. The failing heart interactome includes 602 statistically significant cross-linked peptide pairs altered in the disease condition. Structural insight into ketone oxidation metabolons, OXPHOS machinery, and nucleotide transporter hybrid-conformations, support mitochondrial remodeling in failing heart while bringing forth new hypotheses for pathological mechanisms. Application of quantitative cross-linking technology in tissue provides molecular-level insight to complex biological systems difficult to model in cell culture, thus providing a valuable resource for study of human diseases.

## Introduction

Mitochondria contain over 1,100 proteins that orchestrate diverse cellular functions, from intermediary metabolism and energy production to signaling and cell death (Pagliarini et al., 2008; Rath et al., 2020). Extensive genomic and proteomic approaches have resulted in a curated compendium of mitochondrial proteins with their sub-organellar localization (Rath *et al*., 2020). Advancements in cryo-electron microscopy, X-ray crystallography and NMR have greatly increased understanding of protein function and have enabled structure visualization of purified mitochondrial proteins and complexes with sizes unimaginable only a few years ago (Bridges et al., 2020; Gu et al., 2019; Qi et al., 2021; Spikes et al., 2020; Tucker and Park, 2019). However, within its native cellular or subcellular environment, protein function is dynamically-regulated by the presence of other proteins, post-translational modifications, substrates, cofactors, and transient conformational changes that are not directly measured with conventional techniques.

Chemical cross-linking combined with mass spectrometry (XL-MS) methods have emerged to provide structural insight from complex samples (Sinz, 2018). A cross-linker approach based on peptide synthesis chemistry to introduce MS cleavable features and affinity tags referred to as Protein Interaction Reporter (PIR) technology (Tang et al., 2005) has enabled interactome-level structural studies with isolated functional mitochondria (Schweppe et al., 2017), intact virus particles (Chavez et al., 2012), live cells (Chavez et al., 2016; Weisbrod et al., 2013), and even whole tissue samples (Chavez et al., 2018), providing unique structural insights on protein interaction landscapes. While identification of *in vivo* protein-protein interactions (PPIs) and conformational features from cross-linked peptides increases knowledge about how proteins function within cells, quantitation of cross-linked peptide level changes during perturbation reveal molecular features that confer functional changes at the systems-level. For instance, quantitation of PIR cross-linked peptide levels in cells using SILAC (Ong et al., 2002) revealed functional differences relevant to acquired resistance to topoisomerase I inhibition therapy (Chavez et al., 2015). Moreover, quantitative PIR applications to cellular pharmacological studies revealed interactome changes that are drug-concentration dependent and mechanism-of-action specific (Chavez et al., 2019a; Chavez *et al*., 2016). Thus, quantitative cross-linking technologies enable visualization of interactome changes in living systems relevant to functional changes that could be informative in pathological comparisons. Recently, PIR technologies were further advanced to include isobaric quantitative capabilities that can enable quantitative interactome studies for systems without need for SILAC (Chavez et al., 2020a), as is employed here for failing heart mitochondrial interactome studies.

Due to the extraordinary energy requirements of the heart, cardiomyocytes contain the highest concentration of mitochondria of any cell in the body across mammalian species (Barth et al., 1992; Brown et al., 2017). Concurrently, mitochondrial dysfunction is a well-known maladaptive mechanism in the progression of heart failure. In this study, the feasibility of applying isobaric quantitative PIR (iqPIR) (Chavez *et al*., 2020a) XL-MS technology to mitochondria from healthy and failing mouse hearts is demonstrated. For the first time, integrated measurements of protein interactions, conformations, and surface accessibility allowing for comparative network analysis was enabled directly in tissue. These advancements provide the unique opportunity to study chronic, organ-level conditions, where cell modeling does not holistically recapitulate the bewildering complexity of disease. Of the total ∼3,800 non-redundant cross-linked peptide pairs detected, 90% of cross-linked peptide pairs are quantified in at least one biological replicate. Statistical analysis revealed 602 cross-linked peptide pairs were significantly altered in failing hearts compared to sham controls, corresponding to altered catalytic pockets, hybrid conformations, and higher-order protein assemblies. This study provides molecular-level insight into basic mitochondrial function, and serves as a systems-level resource for generating new hypotheses for disease progression.

## Results

### Quantitation of mitochondrial protein interactome in failing hearts

A workflow was developed to assess mitochondrial protein interactome in failing hearts using iqPIR technology (Fig 1A). Heart failure was induced by transverse aortic constriction (TAC, n=6) surgery in mice. Sham-operated (Sham) animals were used as controls (n=6). TAC hearts demonstrated a decline in left ventricular fractional shortening (Fig 1B) and increased chamber size (Fig S1A-B) compared to Sham. At harvest, TAC mice showed significant cardiac hypertrophy (Fig 1C) and clear signs of pulmonary congestion (Fig S1C-D), indicating the development of heart failure.

**Figure 1:**
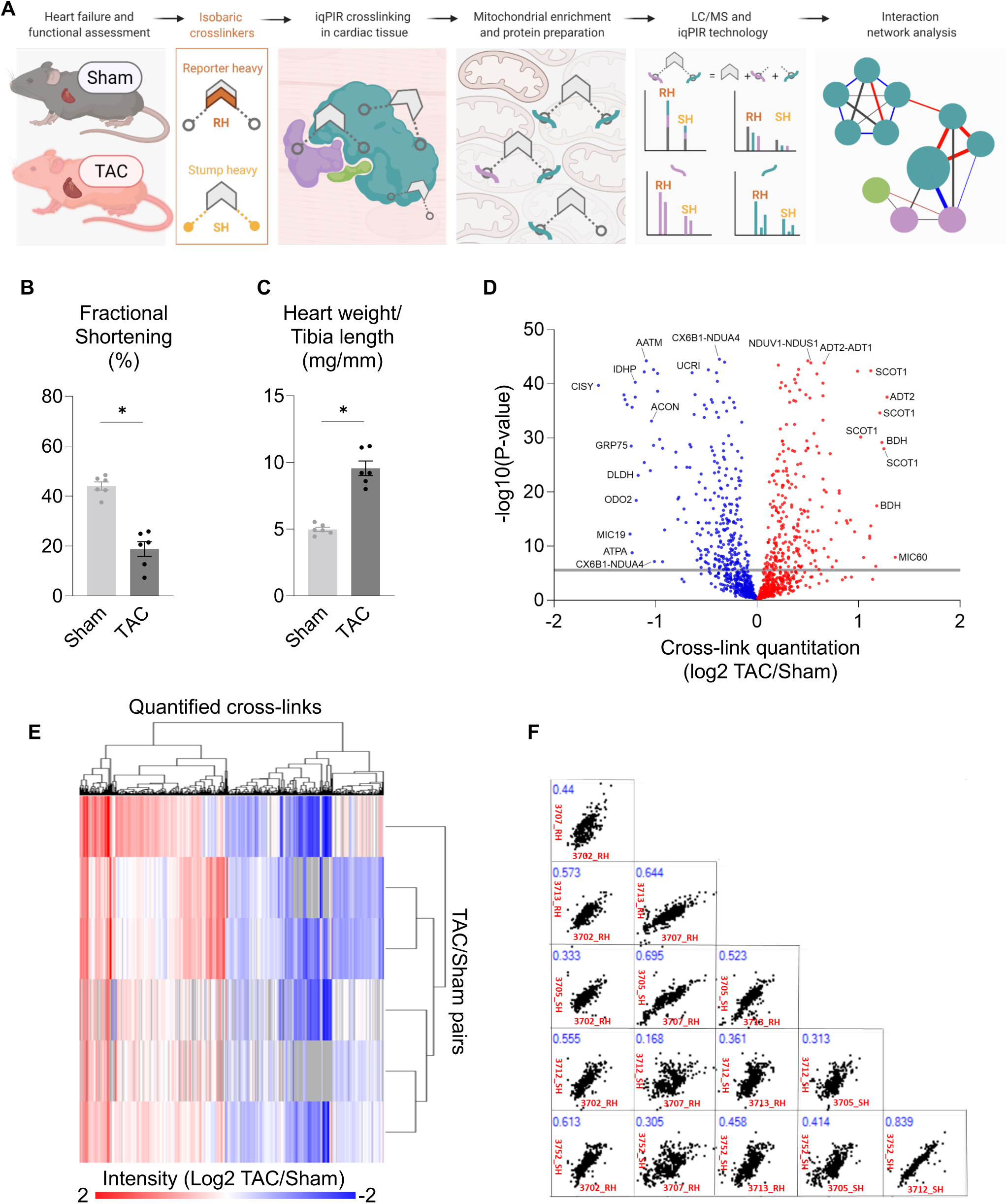
Quantitation of mitochondrial protein interactome in failing hearts. (A) Schematic of quantitative failing heart interactome pipeline. Briefly, TAC and Sham hearts were excised and subjected to stump-heavy and reporter-heavy iqPIR cross-linking reaction followed by mitochondrial enrichment. Samples were pooled for downstream processing and data acquisition by LC/MS. Identical cross-linked peptide pairs from TAC and Sham samples have identical masses yet produce distinct quantitative isotope signatures in MS^2^ spectra, and their intensities enable relative quantitation of cross-linked peptides. Illustration created with BioRender.com. (B) Assessment of cardiac function measured by fractional shortening (%) using echocardiography in TAC and Sham groups four-weeks post-surgery. (C) Cardiac hypertrophy measured by ratio of heart weight (mg) to tibia length (mm) harvest. (D) Volcano plot of quantified cross-link lysine ratios (Log2 TAC/Sham) versus statistical significance. Significance threshold set to p=2.5×10-6 (-log10=5.6). Blue circles indicate cross-linked peptide ratios with decreasing quantitation, while red circles indicate cross-linked peptide ratios with increasing quantitation (relative to TAC). (E) Heat map illustration of quantified cross-linked peptide pairs showing significant changes between TAC and Sham groups, filtered for values present in at least 4/6 biological replicates. (F) Pair-wise scatter plots of 6 pairs of biological replicates. Correlation of Log2 ratios of two biological replicates as shown in X-axis and Y-axis was evaluated with linear regression. R-squared value marked in blue font on top left of each figure. For (B-C), all data are n=6, AVG+/-SEM, * denotes p<0.05 by Student’s t test.

Dissected TAC or Sham cardiac tissue was cross-linked with 2-plex iqPIR cross-linker with either Reporter Heavy (RH) or Stump Heavy (SH) iqPIR versions. TAC and Sham cross-linked samples were then mixed pairwise in a 1:1 ratio based on equal amount of total proteins. Thus, a total of 6 pairs of biological replicates were generated including 3 pairs of forward samples (RH labeled TAC combined with SH labeled sham) and 3 pairs of reverse samples (SH labeled TAC combined with RH labeled sham). The forward/reverse labeling strategy was used to evaluate if there was any quantitation bias caused by labeling direction.

In total, 3,792 non-redundant cross-linked peptide pairs were identified from 6 pairs of TAC/Sham heart samples at an estimated FDR of 1% or less, corresponding to 2,734 lysine residue pairs and 507 protein pairs. The failing heart interactome network is available online in XLinkDB (Schweppe et al., 2016) (xlinkdb.gs.washington.edu, network table name = CaudalCellCommunity2021_Bruce). From the total coverage, over 90% (3607/3792) of the cross-linked peptide pairs were quantified in at least one of 6 pairs of biological replicates. All quantified cross-link ratios were plotted against the p-value of each ratio (Figure 1D), which determined statistical outliers (significance threshold p-value= 2.5×10^-6^). The data were further processed by applying a statistical t-test and Bonferroni correction to all quantified cross-linked peptides, then further filtered to a maximum of 2 missing ratios out of 6 pairs of biological replicates.

A final number of 602 cross-links were quantified showing statistically significant differences between TAC and Sham hearts (Supplemental Table 1). The heat map containing 602 cross-links was generated using NG-CHM BUILDER (Broom et al., 2017) with Euclidean distance and complete linkage selected for hierarchical clustering (Figure 1E). A total of 316 cross-linked peptides were decreased while 286 cross-linked peptides were increased in TAC. iqPIR cross-linkers employ reactive esters to covalently react with surface accessible lysine residues. Dead-end (DE) labeled peptides are generated when only one ester reacts with a lysine residue, while the other ester is hydrolyzed. This feature is useful as a read-out for the surface accessibility of proteins, and an indirect survey of relative protein level changes. Since PPI levels can be affected by endogenous protein levels, DE-labeled peptide quantitation and intra-cross-linked peptides were used to estimate protein abundance for each cross-linked peptide pair as demonstrated previously by Chavez et al (Chavez *et al*., 2020a). For each non-redundant cross-link pair, the mean Log2 ratios for Protein A and Protein B were calculated from all DE peptides (Supplemental Table 1, Fig S1E). In addition, reproducibility of the data was evaluated by applying linear regression to 6 pairs of biological replicates on a pair-wise basis. R-squared values were calculated and shown in Figure 1F. Despite the known individual variations in the TAC model, strikingly high data correlation and consistency were observed across all 15 comparisons of 6 TAC/Sham pairs. It is worth noting that each sample is 1:1 mix of cross-linked proteins from two different animals thus regression fitting of two samples involved 4 different animals. The correlations of forward and forward labeling samples (average R^2^=0.54) are similar to those of forward and reverse labeling samples (average R^2^=0.45) (Supplemental Table 2), which further confirmed the consistency and reproducibility of iqPIR quantitation results. The subset of cross-linked peptide pairs displaying statistically different ratios in TAC were further analyzed by interaction network analysis (Fig S1F). Comprehensive views of pathways and proteins exhibiting large statistical changes (Log 2 ratios are generally ≥1 or ≤-1) led to key highlights of TAC-induced structural changes that will be further discussed in greater detail in the following sections.

### Active conformational states of ketone oxidation proteins enriched in TAC

Recent studies demonstrated an increase in ketone consumption by failing hearts (Aubert et al., 2016; Bedi et al., 2016; Murashige et al., 2020). Enzymes for ketone oxidation, D-β-hydroxybutyrate dehydrogenase (BDH) and Succinyl-CoA:3-oxoacid-CoA transferase (SCOT1), were upregulated in the failing heart but mechanisms driving the flux of ketone oxidation were not fully understood (Aubert *et al*., 2016; Bedi *et al*., 2016; Kolwicz et al., 2016). Circulating ketones, primarily generated by the liver, are taken up by cardiomyocytes and metabolized via three consecutive reactions catalyzed by BDH, SCOT1 and Acetyl-CoA acetyltransferase (THIL) to produce acetyl-CoA for further oxidation in the TCA cycle (Fig 2A). Consistent with increased expression of ketone metabolism proteins in failing myocardium (Aubert *et al*., 2016; Bedi *et al*., 2016), DE quantitation of BDH, SCOT1 and THIL were increased in the iqPIR dataset (Supplemental Table 1). However, quantification of cross-linked lysine residues in SCOT1 and THIL did not match the changes of DE suggesting the presence of protein structural changes beyond changes to abundance (Fig 2B).

**Figure 2:**
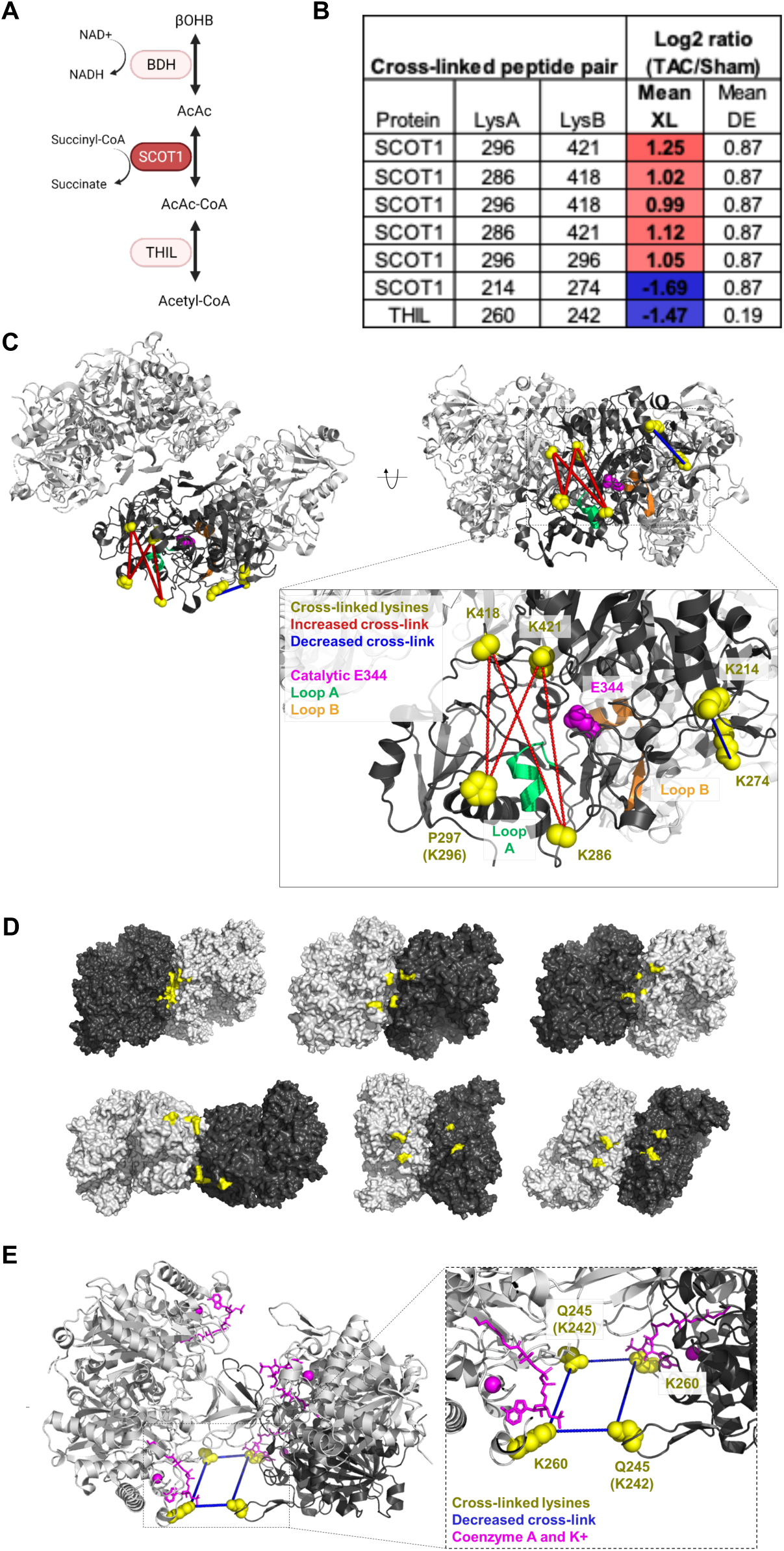
Active conformational states of ketone oxidation proteins enriched in TAC. (A) Schematic of ketone oxidation pathway. (B) Table summarizing the mean cross-linking ratio and DE ratio for cross-linked peptide pairs exhibiting quantitative changes in ketone oxidation machinery. (C) Human SCOT1 tetrameric structure (PDB: 3DLX) shown in grey. Monomeric unit is shown in black centered around the catalytic E344 (magenta). Cross-linked lysine side-chains (K286, K296/P297, K418, and K421, shown as yellow spheres) surround Loop A and together form four cross-links found to increase in TAC. Loop A (CoA binding site, residues 321-329, chain shown in green) is a dynamic region that must undergo conformational change to prevent steric clashing of N- and C-termini during catalysis. Decreasing cross-linked peptide pair, K214-K274, is also shown (yellow spheres) near Loop B (Succinate/Acetoacetate binding site, residues 374-386 chain shown in orange). P297 is shown due to missing density of K296. (D) Six solutions (from top ten highest scoring) from molecular docking of two human SCOT1 tetramers (grey and black, PDB: 3DLX) using Symmdock (Schneidman-Duhovny *et al*., 2005) indicative of higher order oligomerization. Residues surrounding K296 (missing density) are shown in yellow, and are consistently docked in close proximity (Cα-Cα<42Å) to each other, providing evidence for K296-K296 cross-link being between two SCOT1 tetramers. (E) Human THIL (PDB: 2IBW) tetrameric structure with bound CoA and KCl (magenta) highlighting the decreased cross-link between K260-K242/Q245 (sidechains shown as yellow spheres). K260 forms the only direct contact with CoA. K242 resides in the anionic loop meant to capture incoming substrate. Mouse K242 is equivalent to human Q245.

SCOT1, encoded by the *Oxct1* gene, is a tetramer of two inter-connected homodimers (Fig 2C). It transfers a Coenzyme A (CoA) moiety from succinyl-CoA to acetoacetate to form acetoacetyl-CoA and succinate (Fig 2A). The reaction involves E344 in the active site, which sits between two binding pockets for succinate/acetoacetate (residues 321-329, Loop A) and CoA (residues 374-386, Loop B), respectively (Tammam et al., 2007). Four non-redundant cross-linked peptide pairs in SCOT1 were increased during TAC (Fig 2B, S2A). Cross-linked peptides containing K286, K296, K418, and K421 were mapped onto the SCOT1 structure (human SCOT1 PDB: 3DLX) and pinpointed a key surface near Loop A (Fig 2C). During catalysis, the C-terminal undergoes a 17 degree angle domain rotation where Loop A must conform in order to alleviate steric clashing with the static N-terminal (Tammam *et al*., 2007). We hypothesize that the rotation renders the cross-linking of the four peptide pairs possible. Thus, increases in the cross-linked peptide pairs suggest a greater fraction of SCOT1 assumes an active conformational state in TAC. As K286 and K296 reside in a relatively disordered region of SCOT1, where detailed structural information is unavailable. However, large conformational change in the C-terminal domain, where the four cross-linked peptide pairs reside, is observed when apo (PDB: 3OXO, chain A) and substrate-bound (PDB: 3OXO, chain E) porcine SCOT1 monomers are aligned (Fig S2B) (Coker et al., 2010; Fraser et al., 2010) supporting the notion that active enzyme assumes distinct conformation in the region. In humans, naturally occurring mutations in Loop A, such as G324E and L327P, result in SCOT1 loss-of-function (Shafqat et al., 2013), indicating the importance of flexible conformations observed in this region.

The homodimer link between SCOT1 K296-K296 increased ∼2-fold in TAC (p-value=2.13E^-13^) indicating increased interaction between two monomeric units of SCOT1 (Fig 2A). However, the K296-K296 distance mapped onto the human tetramer structure (PDB:3DLX) (Figure S2C) is∼66Å, which exceeds the span possible by iqPIR cross-linkers (Cα-Cα<42Å). These results suggested that SCOT1 exists in an alternative conformation or higher order assembly which is increased in TAC. To determine if this is the case, human SCOT1 tetramers were subjected to molecular docking (Schneidman-Duhovny et al., 2005). With no distant constraints, docking results from 6 of the top 10 solutions (Fig 2D, S3A) consistently placed this region at the interface between two tetramers. The N-terminal domain is relatively static (Fig S2B) during catalysis, yet the cross-linked peptide pair K214-K274 exhibited decreasing quantitation during TAC (Fig 2C). Higher-order docking solutions of SCOT1 place K214 and K274 buried at the oligomeric interface between two tetramers in three solutions (Fig S3B). In these conformations, either K214 or K274 would be inaccessible to cross-linking, which further supports possible enrichment of SCOT1 oligomer in TAC.

In contrast to SCOT1, the cross-link between K242-K260 in THIL decreased in TAC despite a moderate increase of DE (mean Log2 ratio=0.19, Fig 2B). This peptide pair was mapped to lysine residues located on adjacent monomers of the human crystal structure (Fig 2E, Substrate-bound THIL PDB 2IBW). Residue K260 (conserved human K263) helps form the CoA binding pocket and provides the only salt-bridge contact with the 3’phosphate moiety (Haapalainen et al., 2007). Post-translational modification (PTM) at either K242 or K260 provides one possible explanation for decreased cross-linking. However, previous reports demonstrated that hyperacylation at K260 suppressed THIL activity (Dittenhafer-Reed et al., 2015; Still et al., 2013) Increased ketone oxidation, as observed in heart failure (Aubert *et al*., 2016; Bedi *et al*., 2016; Murashige *et al*., 2020), would likely increase salt-bridge occupancy by forming the K260-CoA interaction necessary for catalysis and consequently, decrease cross-linking accessibility (Figure S3C).

Taken together, quantitative iqPIR analysis confirmed the upregulation of ketone metabolism proteins as previously reported, and furthermore, suggest possible mechanisms for active protein conformations in the failing hearts. These findings provide greater insight on how ketone metabolism is potentially adapted in the failing heart that could enable future therapeutic developments.

### Decreased interaction between NDUA4 and C6XB1 affects CIV activity in TAC

Mitochondrial respiration, assessed by oxygen consumption rate (OCR), was reduced in TAC hearts (Fig 3A-B). Multiple mechanisms have been proposed for impaired OCR, among which altered function of electron transport chain (ETC) has been proposed (Brown *et al*., 2017). Quantitative mitochondrial interactome data analysis revealed five cross-linked lysine pairs between NDUA4 and CX6B1 (human COX6B isoform 1) in Complex IV that were significantly decreased in TAC (Fig 3C, Supplemental Table 3). Complex IV (Cytochrome C oxidase, CIV) is the terminal end of the electron transport chain, which accepts two electrons from Cytochrome C (CYC) to reduce oxygen into water. Due to its high sensitivity to detergent conditions (Balsa et al., 2012; Zong et al., 2018) interaction between NDUA4 and other subunits of Complex IV has been difficult to assess. Since iqPIR cross-linkers were applied directly to heart tissue to secure PPIs with covalent bonds, the disruption of detergent-sensitive interactions such as those of NDUA4 was avoided with this approach. Identification of multiple NDUA4 inter-protein cross-linked peptide pairs provides direct and definitive evidence that NDUA4 exists as a component of CIV in cardiac mitochondria and provides insight into the NDUA4-CX6B1 interaction. Cross-linked peptides containing residues K10, K13, and K85 of CX6B1 and K56, K74 and K76 of NDUA4 were mapped to disordered regions of both proteins residing in the IMS (Fig 3D, PDB: 5Z62) making further structural analysis challenging.

**Figure 3:**
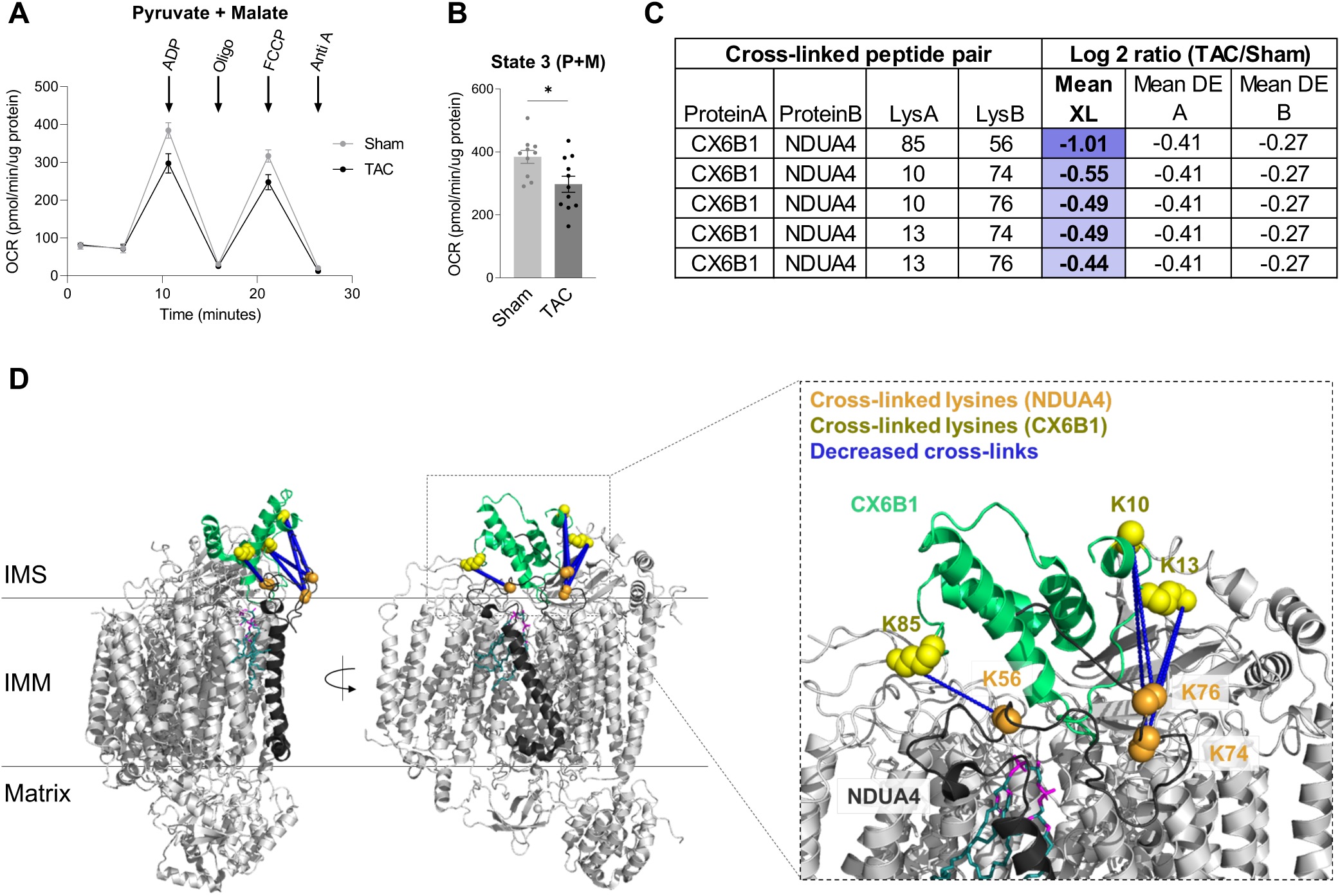
Decreased interaction between NDUA4 and C6XB1 affects CIV activity in TAC. (A) Oxygen consumption rate of mitochondria isolated from TAC and Sham hearts given pyruvate/malate substrates followed by sequential injections of ADP, Oligomycin, FCCP, and Antimycin A, measured by Seahorse XFe24 Analyzer. (B) State III-driven respiration of isolated mitochondria (after addition of ADP). (C) Table summarizing the mean cross-linking ratio and DE level ratio for cross-linked peptide pairs exhibiting quantitative changes in NDUA4 and CX6B1 subunits of CIV. (D) Monomeric Complex IV (grey, PDB: 5Z62). Interactions between NDUA4 (black) and CX6B1 (green) subunits are shown. Cross-linked lysine sidechains are shown as yellow (CX6B1) or orange (NDUA4) spheres. Cross-links exhibit a structural scaffold on the IMS-facing interface of CIV, which are decreased in TAC. Cardiolipin bound to CIV is shown in teal. For (A-B), all data are n=10-11, AVG+/-SEM, *denotes p<0.05 by Student’s t test.

Mutations in CX6B1 R20 which disrupt a salt-bridge with D17 (Massa et al., 2008) are hypothesized to cause instability of CIV leading to encephalomyopathy and hypertrophic cardiomyopathy in patients (Abdulhag et al., 2015). R20 is likely stabilized by a salt-bridge with nearby NDUA4 D60, although this side-chain is only partially resolved in the structure (Figure S4A, PDB: 5Z62). However, Complex IV activity was found to be significantly decreased in TAC (Figure S4B) raising the possibility that NDUA4-CX6B1 interaction modulated CIV function.

Furthermore, as NDUA4 blocks the CIV dimerization interface, its presence maintains CIV primarily as a monomer thus facilitating Complex I-III_2_-IV respirasome assembly (Zong *et al*., 2018). Alterations in the interaction between NDUA4 and other subunits of CIV could therefore affect respirasome formation. Decreased CI-CIII_2_-CIV respirasome has been previously reported in heart failure (Rosca et al., 2008). Thus, quantitative mitochondrial interactome analysis reveals remodeling of Complex IV structure as a new contributor to impaired mitochondrial respiration within the failing heart.

### Enrichment of an intermediate state of ADP/ATP carrier detected in TAC

Exchange of ATP and ADP between mitochondria and cytosol is achieved by the mitochondrial ADP/ATP carrier (ADT), a highly conserved, abundant, and extensively studied translocase (Pebay-Peyroula et al., 2003). ADT is a dual gated transporter that interconverts between at least two known distinct structural conformations (Fig 4A). During a normal cycle, the cytoplasm-open state (C-state) faces the IMS and the gate to the mitochondrial matrix is closed via a series of salt-bridges, allowing for the release of a bound ATP and the binding of a new cytosolic ADP (Pebay-Peyroula *et al*., 2003). The binding of ADP induces a conformational shift to the matrix-open state (M-state) in which the gate to the IMS is closed by formation of new salt-bridges and a hydrophobic plug (Ruprecht et al., 2019). The M-state releases cytosolic ADP in exchange for a newly charged ATP and the cycle repeats. These conformational changes permit the transport of large nucleotide solutes across small protein carriers while preventing proton leak throughout the exchange process (Karch et al., 2019). Mammalian ADT exists in four isoforms which exhibit tissue-dependent but overlapping expression patterns.

**Figure 4:**
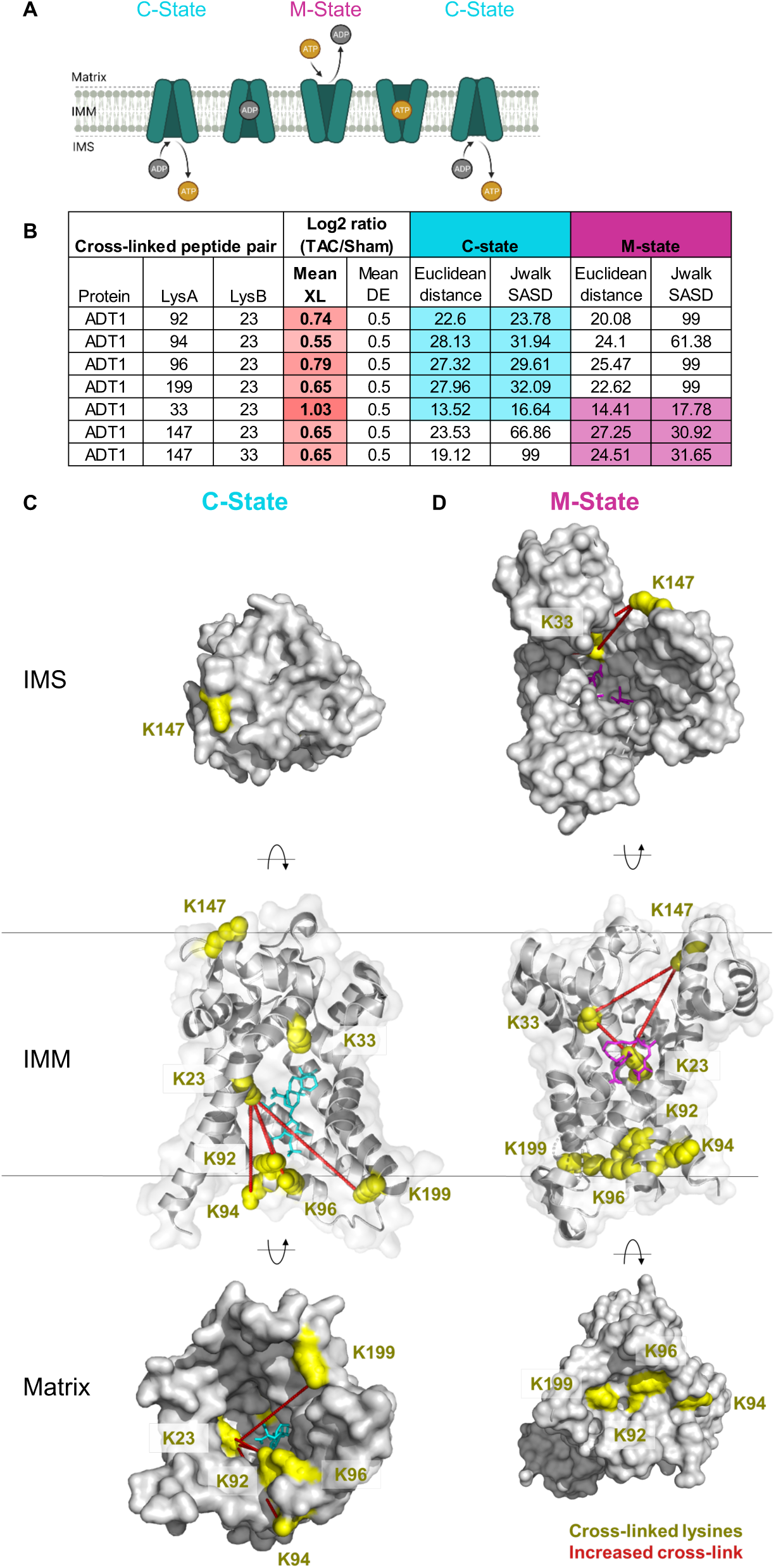
Enrichment of an intermediate state of ADP/ATP carrier detected in TAC. (A) Schematic of ADT cycling between two dominant conformational states. The cytoplasmic-facing state (C-state) delivers ATP to the cytosol and binds ADP. Binding of ADP induces a conformational change to the matrix-facing state (M-state), which returns ADP to the mitochondria and binds a newly charged ATP. (B) Table summarizing mean cross-linking ratio and DE level ratio for each cross-linked peptide pair detected in ADT1 exhibiting changes in TAC. Euclidean distance (EI) and Solvent Accessible Surface distance (Jwalk SASD) (Bullock et al., 2016) are listed for each cross-linked peptide pair when mapped onto both C-state and M-state of ADT1. Highlighted values satisfy iqPIR molecular distance constraint (EI<42Å, Jwalk SASD<51Å). (C) Cross-linked peptide pairs (yellow lysine side chains) mapped onto the C-state conformation of bovine ADT1 (PDB: 1OKC). In this conformation, K23, K92, K94, K96, and K199 are available and can satisfy cross-linking distance constraints. K147 and K33 are not accessible in the M-state. Carboxyatractyloside (CATR) stabilizes the C-state and is shown in cyan. (D) Cross-linked peptide pairs (yellow lysine side chains) mapped onto the M-state of ADT1 (TtAac PDB: 6GCI) where residues K92, K94, K96 and K199 form the gate closed to the matrix, and are inaccessible for cross-linking.Three cross-links are possible and shown between K23-K33, K23-K147, and K33-K147. Bongkrekic acid (BKA) stabilizes the M-state and is shown in magenta.

Quantitative iqPIR analysis revealed increased levels of seven non-redundant cross-linked peptide pairs of ADT1 in TAC (Fig 4B, S4C). It should be noted that other ADT isoforms and multimeric ADT structures may also be present that could explain increases in these observed links (Figure S4D), but ADT1 is the predominant isoform in cardiac and skeletal muscle (Bround et al., 2020). Cross-links between K23-K92, K23-K94, K23-K96 and K23-K199 are compatible with the C-state (Pebay-Peyroula *et al*., 2003) of ADT1 because all lysine residues are solvent exposed in this conformation (Fig 4C, S4B, bovine ADT1 PDB: 1OKC, **Supplemental Video 1**). However, these links are not compatible with the M-state because K92, K94, K96, and K199 must penetrate occluded volume of the protein in this conformation.

Moreover, K96 and K199 form salt bridge interactions with D196 and D292 that contribute to the ADT1 gate closure to the IMS (Fig 4D, S4E, TtAac ADT1 PDB: 6GCI, Supplemental Video 1). Although the K23 and K33 cross-link is compatible with distance constraints, it is known that K33 partakes in the hydrogen-bond network needed to stabilize the C-state by forming a salt-bridge with D232 (Pebay-Peyroula *et al*., 2003) (Fig S4F). On the other hand, cross-links between K23-K33, K23-K147, and K33-K147 are compatible only with the M-state (Fig 4D, S4D). Thus, increases of all seven links would indicate that both C-state and M-state have increased in TAC. This is, however, unlikely given the mutual exclusivity of M- and C-states and the observation of no large change in ADT1 protein levels (Log2 TAC/Sham= 0.5). Functionally, increased enrichment of C-state in failing hearts, in which ATP production is known to be impaired, is also counter-intuitive. Collectively, increases in these seven cross-linked peptide pairs and ADT1 protein level measurements in TAC hearts cannot be explained by either a shift between C-state and M-state of ADT1 in heart failure.

Perhaps a more likely explanation is the existence of an alternative state of ADT1, with which all quantified cross-links are compatible, and this new conformation has increased in TAC. To satisfy all the cross-links in one single state, lysine residues involved in gating at either C- or M-state must be accessible to cross-linkers. This new conformation could be consistent with the P-state which was recently proposed to be simultaneously open to both the IMS and the matrix, synonymous with non-selective mPTP conductance (Karch *et al*., 2019; Ruprecht *et al*., 2019). Although there is currently no known structural evidence, the non-selective P-state is hypothesized to provide an open channel where the lysine residues responsible for gating are not involved in salt bridge formation and thus, may be available for cross-linking. Thus, our results provide novel structural evidence that heart failure is associated with increased conformational enrichment of ADT1 that is non-functional for ADP/ATP translocation but likely possesses non-selective conductivity (Bertholet et al., 2019).

## Discussion

In this study, iqPIR technology achieved one of the highest levels of cross-linking complexity resolved from tissue thus far. Our comparative studies of mitochondria in normal and failing hearts yield quantitative changes in 98 mitochondrial proteins, providing a system’s view of protein structure remodeling in heart failure. Among the 602 cross-linked peptide pairs that showed statistically significant level differences in TAC vs. Sham hearts, enrichment of active ketone oxidation enzymes, altered interactions among Complex IV subunits and novel conformational enrichment of the ADP/ATP transporter were revealed in TAC hearts. These findings advance mitochondrial and heart failure research in several ways. First, the data elucidate new structural information on key players in mitochondrial function, providing a basis for developing future therapies. Second, the results enable generation of novel hypotheses for mechanisms of mitochondrial maladaptation in disease. Most importantly, the quantitative interactome dataset provides a valuable resource for exploration and visualization of changes in many other proteins not discussed here that may be important for improved understanding of mitochondrial function and heart failure pathology.

Using iqPIR technology as a discovery-based approach in this study, conformational and interaction changes in mitochondrial proteins and complexes were observed that possess direct functional relevance to metabolic changes in heart failure. Identification of SCOT1 oligomer metabolons and increased active conformational states of SCOT1 and THIL in the present study are corroborated by previously reported metabolic changes in failing hearts.

Therefore, this agreement serves as affirmation of the utility of quantitative *in vivo* cross-linking and mass spectrometry as a means to gain greater molecular-level insight on changes in heart failure. Importantly, changes in ketone metabolism has been thus far attributed to substrate and enzyme abundance in heart failure with no information on regulatory mechanisms (Kolwicz *et al*., 2016). Our results provide new insight into metabolic remodeling, suggesting that active conformations exist beyond mere increases in protein levels.

The first direct evidence for the NDUA4 existence as a CIV subunit within cardiac tissue was provided by PIR technology (Chavez *et al*., 2018). The present iqPIR data further confirms that interaction and enables the first quantitation of the NDUA-CX6B1 interaction in failing and control heart tissues with five non-redundant Lys-Lys residue pairs between NDUA4 and CX6B1. The decrease in NDUA4-CX6B1 interactions and concomittant decrease in CIV activity in heart failure samples suggests this interaction could serve as a potential modulator of CIV activity. NDUA4 is thought to maintain CIV monomeric population by occupying the CIV dimerization interface. Decreased NDUA4-CX6B1 cross-links, coincident with decreased CIV activity in TAC, supports the hypothesis that monomer and respirasome populations are functionally active populations (Zong *et al*., 2018). It remains unclear whether destabilization of NDUA4 during TAC results in consequential alterations of CIV dimeric pools, and what role this population, if any, may have in disease pathology (Ramzan et al., 2019). Nonetheless, the present observation that NDUA4-CX6B1 interactions decrease in heart failure presents a new, previously unrecognized target for future studies and possible therapies that stabilize this interaction as treatment to restore or prevent CIV functional decline in heart failure.

Mitochondrial ADP/ATP carrier (ADT) has long been proposed to contribute a pore-forming component of the mitochondrial permeability transition pore (mPTP) (Karch *et al*., 2019; Kokoszka et al., 2004). However, several decades of genetics and physiological studies have yielded ambiguous evidence. Transient opening of mPTP has physiological function such as regulating mitochondrial calcium homeostasis or reactive oxygen species signaling while prolonged mPTP contributes to cell death under stress conditions including heart failure (Bround *et al*., 2020). Our results provide critical structural information consistent with recently proposed P state of ADT which opens to both matrix and inter-membrane space (Bround *et al*., 2020). It remains unclear whether the P-state is static, or is resultant from rapid interconversion between two incompletely gated structures. Nonetheless, increased presence of such a state compromises nucleotide translocation function and increases mitochondrial proton leaks leading to impaired energetics. In support of this notion, a mitochondrial-targeted tetrapeptide, elamipretide (SS-31), is shown to bind K23 of ADT1 and reduces proton leak in mitochondria of aging animals (Chavez et al., 2020b). Thus, changes in the conformational state of ADT warrants further investigation both as a disease mechanism and a therapeutic target.

In summary, we demonstrate that iqPIR and mass spectrometry constitute a powerful technology to quantify protein structural changes at the systems-level in tissues. Application of this technology to animal models of heart failure generated a new mitochondrial protein interactome that revealed novel mechanisms that underpin potential therapies.

## Acknowledgments

We thank all members of the Tian and Bruce labs their thoughtful discussion and support. We thank the University of Washington Proteomics Resource for advice and helpful discussions. We thank Dr. Yun-Wei A. Hsu for assistance with the animal models. We thank Dr. Julia Ritterhoff and Dr. Frauke Drees for technical support and guidance.

## Sources of funding

This work was supported in part by U.S. National Institutes of Health (NIH) grants HL110349, HL129510, HL142628 (to R.T.), HL144778, GM097112, R35GM136255 (to J.E.B.), American Heart Association Predoctoral Fellowship 20PRE35120126 (to A.C.), AHA Postdoctoral Fellowship 18POST33990352 (to B.Z.), and NIH 2T32DK007247-41 to (M.A.W).

## Contributions

A.C., X.T., J.D.C., R.T., and J.E.B designed the experiments. A.C., X.T., J.D.C, A.K, M.A.W, R.T., and J.E.B. wrote the manuscript. A.C, X.T, R.T, and J.E.B edited the manuscript. A.C., X.T., J.D.C., A.K., performed formal analysis. A.C., B.Z., M.A.W performed animal experiments. A.C. and J.D.C., performed cross-linking experiments. J.D.C performed protein preparation. J.D.C, X.T., and A.K performed mass spectrometry raw data acquisition and processing. A.K. developed computational tools to support structural protein analysis and cross-linking quantitation. O.V. performed TAC surgery. R.T and J.E.B. supervised the project.

## Declaration of interests

None declared.

---

**Supplemental Figure 1.**
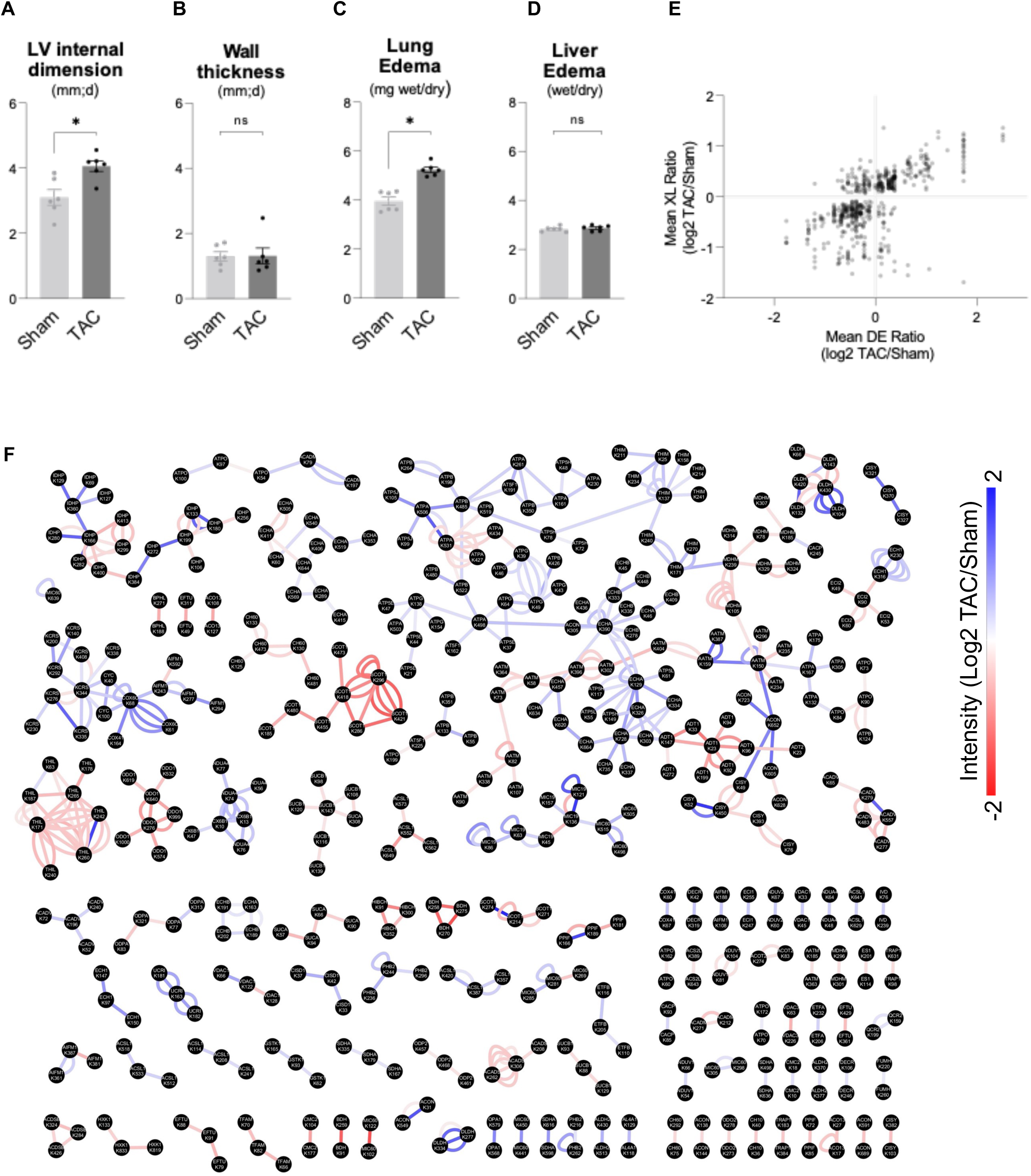
(A-B) Left ventricle internal dimension and wall thickness in TAC and Sham groups determined by echocardiography four weeks post-surgery. (C-D) Lung and liver edema (wet weight/dry weight in mg) measured at tissue harvest. (E) Quantitation of mean cross-link (XL) ratio vs mean dead-end (DE) ratio for each cross-linked peptide pair statistically changed in TAC in at least 4/6 biological pairs (Log2 TAC/Sham). Sum of Mean DE ratio for Protein A and Protein B is shown to account for cross-links between two different proteins. (F) Interaction network of lysine residues (black nodes) connected by observed cross-links (edges). Edges are colored according to increasing (red) and decreasing (blue) quantitation of statistically significant subset of cross-links (Log2 TAC/Sham) shown by color scale. For (A-D), all data are n=6, AVG+/-SEM, * denotes p<0.05 by Student’s t test.

**Supplemental Figure 2.**
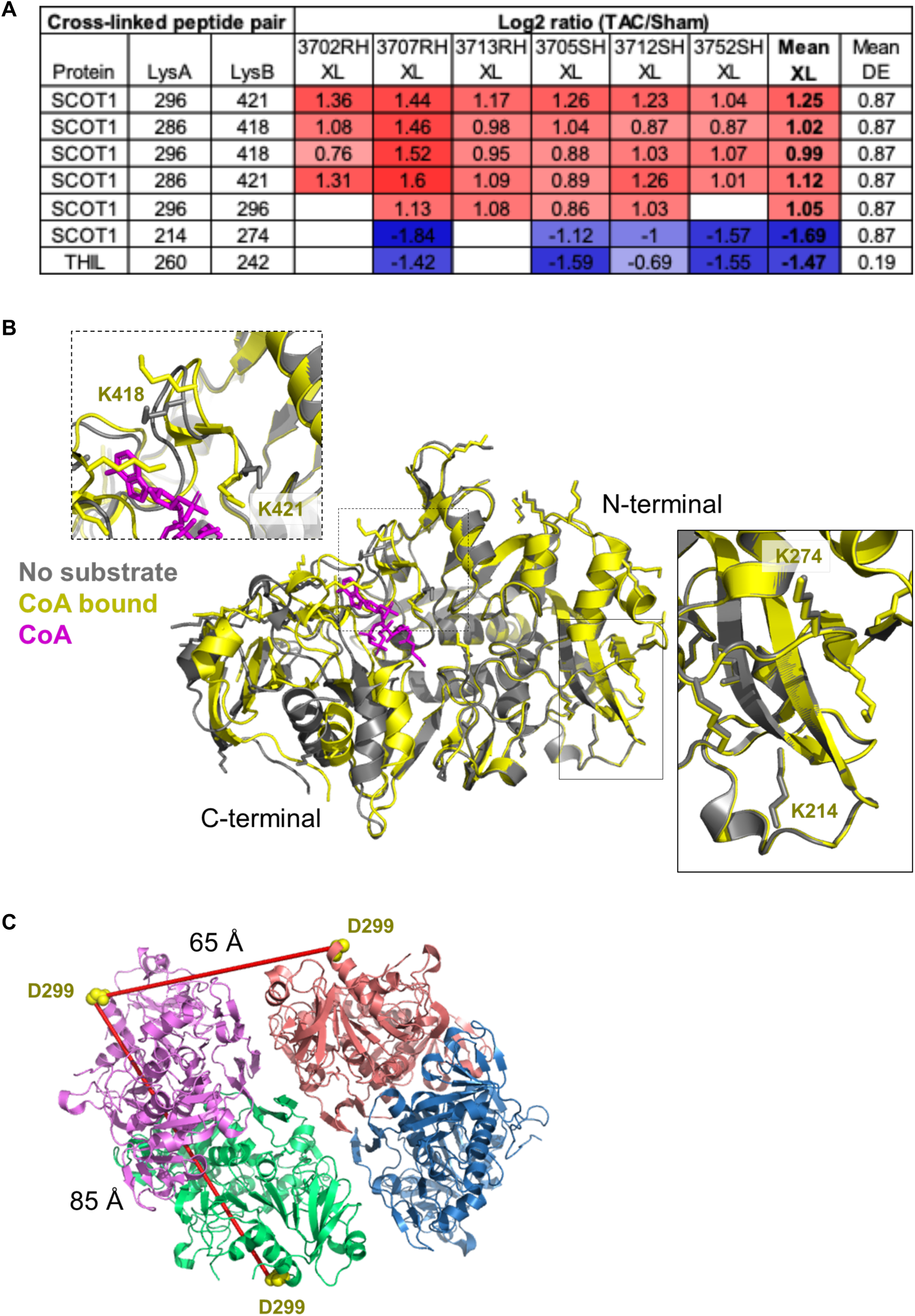
(A) Table summarizing the mean cross-linking ratio and DE ratio for cross-linked peptide pairs including values obtained across biological replicates for ketone oxidation proteins. (B) Alignment of apo (grey, PDB: 3OXO chain A) and substrate-bound (yellow, PDB: 3OXO chain E) monomers of porcine SCOT1. CoA is colored in magenta and bound to the active site. Lysine sidechains are shown in stick representation. Alignment depicts the structural differences between the dynamic C-terminal domain and the static N-terminal domain during substrate-binding. Close up view specifies cross-linked lysines (K214, K274, K418, and K421). (C) Human SCOT1 tetrameric structure (PDB: 3DLX) illustrating the observed K296-K296 homodimer cross-link that increases in TAC. Molecular interaction distance constraint is exceeded when mapped onto the tetrameric structure. Each monomer is depicted in a different color. K296 is not present in electron density of solved structures, closest resolved residue present in each monomer is shown (D299).

**Supplemental Figure 3.**
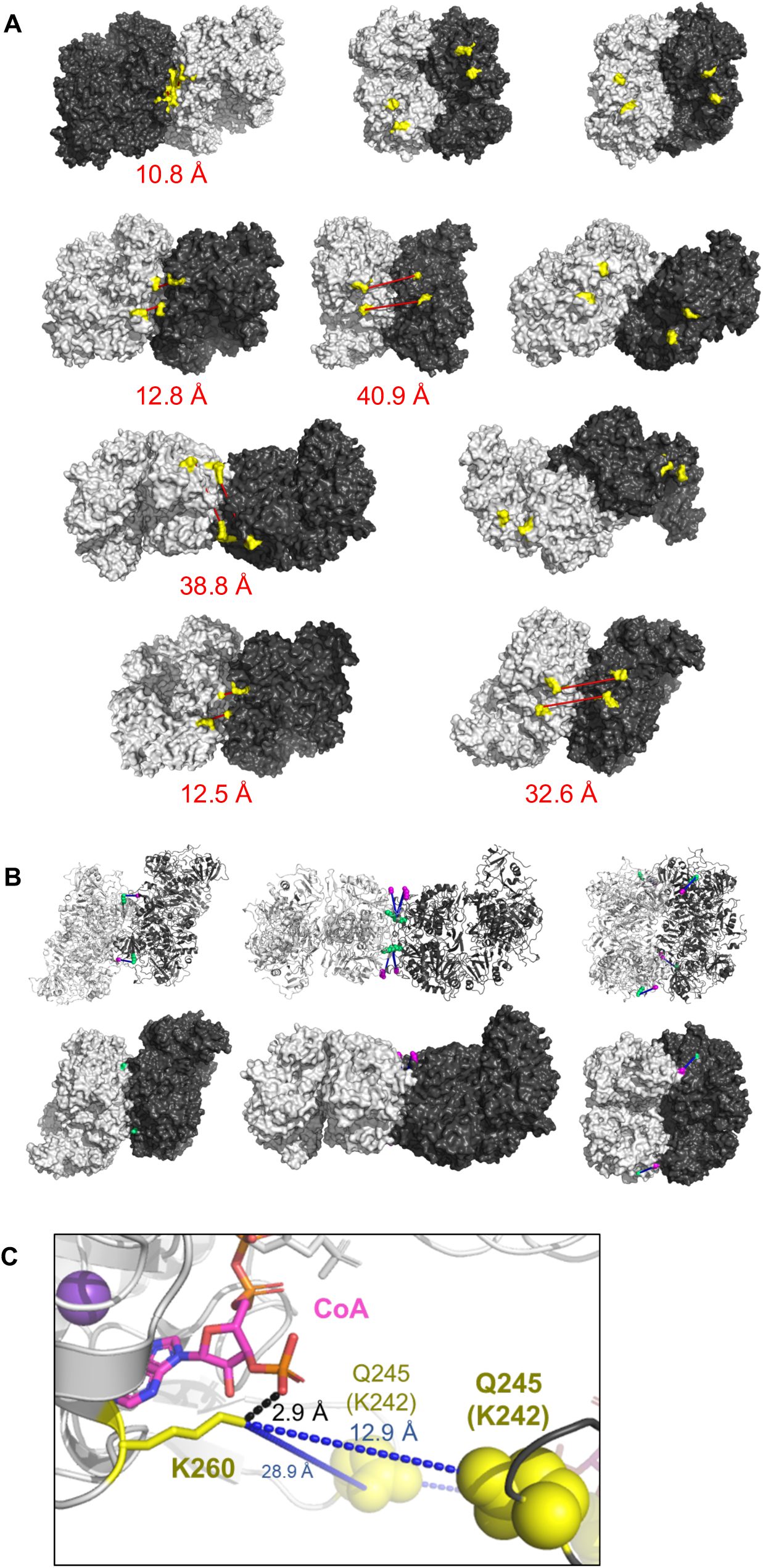
(A) Top 10 solutions from molecular docking of two human SCOT1 tetramers (PDB: 3DLX) using Symmdock (Schneidman-Duhovny *et al*., 2005). Approximate location of K296-K296 was examined in each solution. Due to the missing density around K296, residues 284-302 were highlighted (yellow). In 6 of the top 10 solutions, proximity between two tetramers is consistent with molecular distance constraint of cross-linker (listed below structure). (B) Three (of top ten) docking solutions where residues K214 or K274 are buried at the interface between two SCOT1 tetramers (PDB: 3DLX). Decreasing cross-link between K214 (green) and K274 (magenta) is shown as a blue edge. (C) Human THIL (PDB: 2IBW) tetrameric structure highlighting the salt bridge formed between K260 (yellow stick representation) and 3’ phosphate moiety of CoA (2.9 Å), which may make K260 inaccessible for cross-linking with an adjacent K242/Q245 (yellow sphere) (intra-link distance 28.9 Å) or K242/KQ245 on the opposing monomer (inter-link distance 12.9 Å).

**Supplemental Figure 4.**
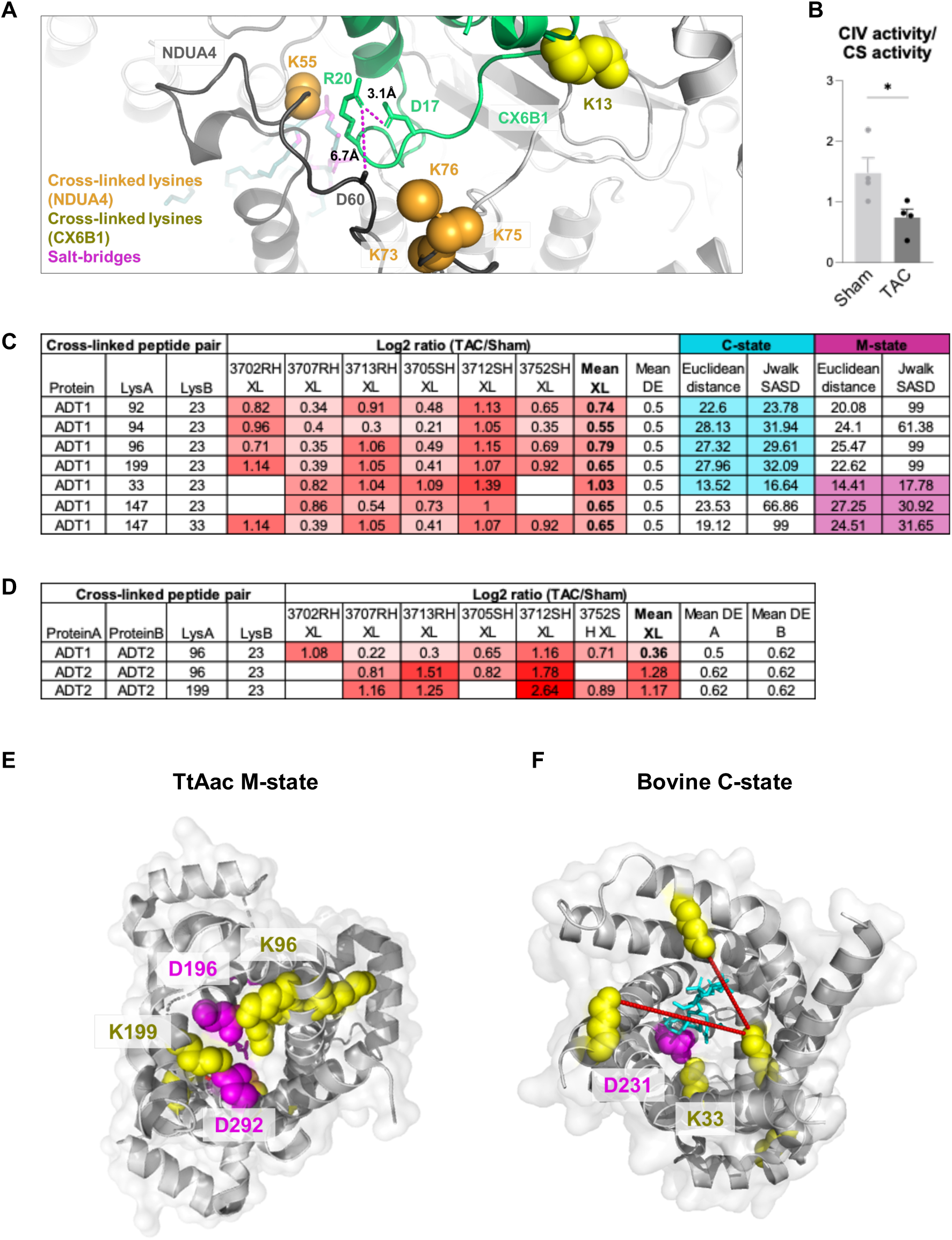
(A) Structural insight into CX6B1 R20 forming salt-bridges at NDUA4-CX6B1 interface. R20 sidechain (green) forms a salt-bridge with CX6B1 D16 (green) and NDUA4 D60 (black), which pinpoints an interface necessary for stability of CIV. Side-chains of R20, D16, and NDUA4 D60 (partially resolved) are depicted in stick representation. Cross-linked lysine sidechains are shown as yellow (CX6B1) or orange (NDUA4) spheres. (B) Cytochrome C oxidase enzymatic activity assay in tissue homogenates from TAC and Sham hearts, normalized to Citrate Synthase activity. N=4, AVG+/-SEM, *denotes p<0.05 by Student’s t test. (C) Table summarizing the mean cross-linking ratio and DE ratio for cross-linked peptide pairs including values obtained across biological replicates for ADT1. (D) Table summarizing the mean cross-linking ratio and DE ratio for cross-linked peptide pairs including values obtained across biological replicates for cross-links in ADT isoforms. (E) Cross-linked peptide pairs (yellow lysine side chains) mapped onto the M-state conformation of ADT1 (TtAac PDB: 6GCI). Salt-bridges between K96-D196 and K199-D292 contribute to the gating mechanism which closes the M-state to the IMS (Ruprecht *et al*., 2019), and would make lysines unavailable for cross-linking. Aspartic acid sidechains are shown in magenta. (F) Cross-linked peptide pairs (yellow lysine side chains) mapped onto the C-state conformation of bovine ADT1 (PDB: 1OKC). K33 and D231 are known to form a salt-bridge which stabilizes the C-state (Pebay-Peyroula *et al*., 2003) and would make K33 unavailable for cross-linking. Aspartic acid sidechains are shown in magenta.

**Supplemental Video 1**

This video can be viewed by anyone with the following link: https://drive.google.com/file/d/1G8NtkRJylq1xKNzMa4ovpwFlupzenzht/view?usp=sharing

## STAR METHODS

### KEY RESOURCES TABLE

| REAGENT or RESOURCE | SOURCE | IDENTIFIER |
| --- | --- | --- |
| **Chemicals, Peptides, and Recombinant proteins** |  |  |
| iqPIR cross-linker | AnaSpec, Chavez et al 2020 | Peptide sequence: Gly-GlyLys(biotin)-Lys-Pro<sub>2</sub>-Asp<sub>2</sub>-succinate<sub>2</sub>-N-hydroxyphthalimide (NHP)<sub>2</sub> |
| **Deposited Data** |  |  |
| LC-MS data deposited into PRIDE | https://www.ebi.ac.uk/pride/archive/ | Accession No: PXD027757 |
| Cross-link data deposited into XLinkDB | http://xlinkdb.gs.washington.edu/xlinkdb/index.php | Network Name: CaudalCellCommunity2021_Bruce |
| **Experimental Models: Organisms/Strains** |  |  |
| Mouse: C57BL6/NCrl | Charles River Laboratories | IMSR_CRL:27 |
| **Software and Algorithms** |  |  |
| Comet v. 2018 | Eng et al 2013 |  |
| Mango | Moir et al 2018 |  |
| Xlink Prophet | Keller et al 2002 |  |

### RESOURCE AVAILABILITY

#### Lead contacts

Further information and requests for resources and reagents should be directed to and will be fulfilled by the lead contacts, Rong Tian and James E. Bruce.

#### Materials Availability

This study did not generate new unique reagents.

#### Data and software availability

Cross-linking data has been deposited at XlinkDB and is publicly available as of the date of publication. DOI is listed in the key resources table. The mass spectrometry proteomics data have been deposited to the ProteomeXchange Consortium via the PRIDE (Perez-Riverol et al., 2018) partner repository with the dataset identifier: PXD027757. Any additional information required to reanalyze the data reported in this paper is available from the lead contacts upon request.

### EXPERIMENTAL MODEL AND SUBJECT DETAILS

#### Animal Model

All protocols concerning animal use were approved by the Institutional Animal Care and Use Committee at University of Washington. This study utilized 12 wild-type mice, strain C67Bl6/NCrl (IMSR_CRL:27). Female and male mice have different responses to pressure overload. TAC surgery causes a mild and variable phenotype in female. Thus, only male mice were used in this study. Adult (>10 weeks old, weighing 24-26g) male mice were chosen randomly into experimental groups. Mice were housed in a vivarium with a 12 hr light/dark cycle at 22°C. Mice were maintained on ad libitum standard rodent diet and water.

### METHOD DETAILS

#### Transverse aortic constriction (TAC) surgery

Male mice aged 10-12 weeks, weighing 24-28g underwent TAC or Sham surgery as previously described (Tarnavski et al., 2004). Mice were anesthetized with 4% isoflurane and intubated with a 20-gauge canula. Mice were ventilated at 2.5% isoflurane at 135 breaths per minute by a small animal TOPO ventilator (Kent Scientific, Torrington, CT). The aortic arch was exposed and separated from the thymus via left thoracotomy. A 27-gauge blunt needle was held near the aorta (between the brachiocephalic and left common carotid arteries). A constriction of the transverse aorta was generated by tying a 6-0 Ethilon ligature against the blunt needle. The needle was then promptly removed. The lungs were inflated and the chest was closed by 5-0 polypropylene suture. The animal was removed from the ventilation system and given SR buprenorphine (subcutaneous, 0.5mg/kg) analgesic and 0.9% saline (intraperitoneal, 0.2ml) for hydration after mice regained consciousness (2hrs after ventilation). Sham operated mice underwent all the same procedures as TAC, excluding ligature of the aorta. Mice were randomly assigned to TAC and Sham procedures, and researcher was blind to operation until after echocardiogram analysis was performed. Combined mortality (acute and chronic) was less than 25%.

#### Transthoracic echocardiography

Mice were anesthetized and maintained with 0.8%-2% isofluorane in 95% oxygen at rates of 550-600 beats per minute as done previously(Ritterhoff et al., 2020). Trans-thoracic echocardiography was conducted 4 weeks post-TAC and Sham surgery with Vevo 2100 high-frequency, high resolution digital imaging system (VisualSonics) with a MS400 Microscan Transducer. A parasternal long-axis view was used to collect M-mode images for analysis of fractional shortening, ejection fraction, and other functional parameters.

#### Cross-linking of cubed heart tissue

Mice were euthanized by cervical dislocation. The chest cavity was opened, and the heart was rapidly excised and placed into ice-cold, mitochondria isolation medium (MIM: 70mM sucrose, 220mM mannitol, 5mM MOPS, 1.6mM carnitine hydrochloride, 1mM EDTA, 0.025% fatty acid-free BSA, pH 7.4 with 5M KOH) to flesh out blood. The aorta and atria were removed. The myocardium was weighed (heart weight). The heart was transferred to a pre-chilled 60mm dry petri dish and maintained on ice. Heart tissue was finely minced with a razor blade to a homogenous tissue size distribution of approximately 1mm^2^ cubes.

Heart tissue was centrifuged at 1500 x g for 5 min at 4°C, and MIM was replaced with cross-linking buffer (170 mM Na2HPO4, pH 8). iqPIR (Chavez *et al*., 2020a) is a novel technology recently developed for quantitation of cross-linked peptide pairs using isobaric stable isotopes selectively incorporated into the PIR cross-linker, which enables MS^2^-based interactome quantitation in a way analogous to how TMT (Thompson et al., 2003) or iTRAQ (Ross et al.) is used for proteome quantitation. Briefly, reporter heavy (RH) or stump heavy (SH) iqPIR cross-linkers were applied to different samples individually first, then TAC/Sham cross-linked sample pairs were pooled together for further downstream sample processing and data acquisition. This strategy greatly decreases sample handling errors and reduces instrument time for data acquisition. Identical cross-linked peptide pairs from TAC and sham samples have identical masses and LC retention times, yet produce different and distinct quantitative isotope signatures in MS^2^ spectra. The intensities of TAC- or Sham-distinct isotope labeled peptide and fragment ions in each MS^2^ spectrum enable relative quantitation of cross-linked peptides in TAC and Sham samples. The samples were mixed for 30 min at room temperature on a Thermomixer at 800 rpm. The tissue was centrifuged at 1500 x g for 5 min at 4°C and the supernatant was removed. Cross-linked tissue was then further subjected to modified mitochondrial isolation as described below.

#### Mitochondrial isolation from cross-linked heart tissue

Cross-linked heart tissue was washed in fresh MIM and centrifuged at 1500 x g for 5 min at 4°C. Subsequent steps are the same as regular mitochondrial isolation protocol below. However, resulting fractions and supernatants were collected after each step. All fractions were frozen in -80°C after isolation and saved for processing.

#### Protein extraction from cross-linked mitochondria

Frozen fractions were transferred to a stainless steel cryogrinding jar cooled to -196°C with liquid nitrogen in 0.1M NH_4_HCO_3_. The samples were cryoground for five 3 min cycles at 30 Hz using a Retch MM 400 mixer mill. Between cycles, the cryogrinder was cooled with liquid nitrogen between cycles. The resulting frozen powder was transferred to a falcon tube where 8M urea (in 0.1M Tris, pH 8.0) was added. Samples were sonicated using a GE-130 ultasonic processor, followed by reduction of cysteine residues by incubation with 5mM Tris (2-carboxyethyl) phosphine (TCEP, Fisher Scientific) for 30 min followed by alkylation with 45 min incubation with 10mM iodoacetamide (Fisher Scientific). Urea concentration was reduced to less than 1M by diluting the samples by a factor of 10 with fresh 0.1M Tris Buffer (pH 8.0). The protein concentration was measured using the Pierce Coomassie protein assay (Thermo Scientific).

#### Protein digestion and enrichment of cross-linked peptides

The procedures for protein digestion, desalting, and enrichment of cross-linked peptides by strong cation exchange chromatography (SCX) and biotin-capture were followed as described in a previous publication (Chavez et al., 2019b). Briefly, extracted protein were digested with trypsin (200:1 in weight ratio) overnight at 37°C and then quenched with acidification of digest to pH ∼3 by adding TFA. The digest was then desalted using Sep-Pak cartridges and fractionated on SCX using a flow rate of 1.5 mL/min and 97.5-min gradient with an increasing percentage of SCX solvent B (solvent A: 7 mM KH_2_PO_4_, pH 2.6, 30% (vol/vol) acetonitrile and solvent B: 7 mM KH_2_PO_4_, pH 2.6, 350 mM KCl, 30% (vol/vol) acetonitrile). The SCX fractions were adjusted to pH 8.0 and then subjected to biotin capture by adding monomeric avidin UltraLink resin (Thermo Scientific) to each pooled fraction. After 3 times of washing the avidin beads with 0.1 M NH_4_HCO_3_, pH 8.0, the cross-linked peptides were eluted with 70% (vol/vol) acetonitrile, 1% (vol/vol) formic acid (FA) and then dried in a speed vacuum. The dried samples were reconstituted in 0.1% FA and ready for LC-MS/MS analysis.

#### LC-MS/MS

Cross-linked peptide samples were analyzed in technical duplicate by liquid chromatography mass spectrometry using an Easy-nLC (Thermo Scientific) coupled to a Q Exactive Plus mass spectrometer (Thermo Scientific). Each analysis used a 3 μL injection of sample onto a 3 cm x 100 μm inner diameter fused silica trap column packed with a stationary phase consisting of 5 μm Reprosil C8 particles with 120 Å pores (Dr. Maisch GmbH) with a flow rate of 2 μL/ min of mobile phase consisting of solvent A (H2O containing 0.1% formic acid) for 10 minutes.

Peptides were then fractionat-ed over a 60 cm x 75 μm inner diameter fused silica analytical column packed with 5 μm Reprosil C8 particles with 120 Å pores by applying a linear gradient from 95% solvent A, 5% solvent B (acetonitrile containing 0.1% formic acid) to 60% solvent A, 40% solvent B over either 120 or 240 minutes at a flow rate of 300 nL/min. Eluting peptide ions were ionized by electrospray ionization by applying a positive 2.2 kV potential to a laser pulled spray tip at the end of the analytical column. The mass spectrometer was operated using a top five data dependent acquisition method with a resolving power setting of 70,000 for MS^1^ and MS^2^ scans. Additional settings include an AGC target value of 1e6 with a maximum ion time of 100 ms for the MS^1^ scans and an AGC value of 5e4 with a maximum ion time of 300 ms for the MS^2^ scans. Charge state exclusion parameters were set to only allow ions with charge states from 4+ to 7+ to be selected for MS^2^. Ions selected for MS^2^ were isolated with a 3 m/z window and fragmented by HCD using a normalized collision energy setting of 30. Ions for which MS^2^ was performed were then dynamically excluded from further selection for MS^2^ for 30 s.

#### Heat Map Analysis

Perseus (version 1.6.15.0) (Tyanova et al., 2016) was used for statistical data analysis of cross-link quantitation ratios. Data was first filtered based on 70% of total valid values across all biological replicates and then applied with T-test using Benjamini-Hochberg FDR at 0.05. T-test q-values less than 0.05 were considered significant. Heatmaps were generated using NG-CHM GUI software (version 2.20.0) (Broom *et al*., 2017) with Euclidean distance and complete linkage selected for both row and column hierarchical clustering.

#### Mitochondrial Isolation from cardiac tissue

Hearts were excised from mice, the aortas and atria were removed. Heart tissues were rinsed briefly in ice-cold mitochondria isolation medium (70mM sucrose, 220mM mannitol, 5mM MOPS, 1.6mM carnitine hydrochloride, 1mM EDTA, 0.025% fatty acid-free BSA, pH 7.4 with 5M KOH) with the addition of 2 mM taurine, to remove residual blood. Tissues were minced on ice and resuspended in fresh MIM, followed by trypsin digestion (10ug/ml) and incubated on ice for 10 min. Trypsin digestion was stopped by the addition of trypsin inhibitor (0.5mg/ml) and additional BSA (1 mg/ml) to MIM. The suspension was centrifuged for 1 min at 1,500 x g at 4°C, and the supernatant was discarded. The tissue pellets were resuspended in fresh MIM containing 1mg/ml BSA, and transferred to a Teflon-glass tube and homogenized on ice with a Teflon pestle. The homogenates were centrifuged for 10 min at 800 x g at 4°C. The supernatants were collected and centrifuged for 10 min at 8,000 x g at 4°C. The supernatant was discarded and the mitochondrial pellets were resuspended in MIM to wash. The resuspension was centrifuged for 10 min at 8,000 x g at 4°C, and the supernatant discarded. The mitochondrial pellet was resuspended to a concentrated volume for protein quantification by BCA Assay (Pierce).

#### Mitochondrial respiration using Seahorse XFe24 Analyzer

The oxygen consumption rate (OCR) of isolated cardiac mitochondria was measured with an XF24e Flux Analyzer (Seahorse Bioscience, Agilent) based off the manufacturer’s protocol(Rogers et al., 2011). The Seahorse utility plate was hydrated with calibration buffer and incubated overnight at 37°C prior to respiration experiments. The mitochondria were seeded in 24-well microplates at 1-2ug protein per well in 1x mitochondrial assay solution (70mM sucrose, 220mM mannitol, 10mM KH_2_PO_4_, 5mM MgCl_2_, 2mM HEPES, 1mM EGTA, 0.2% BSA, pH 7.35 at room temperature) containing substrates and centrifuged for 2,000 x g for 20 min at 4°C. Respiration buffer was added to each well to a final volume of 500ul, then the plate was placed in a non-CO2 incubator for 10 min at 37°C. The plate was incubated the Seahorse Analyzer and a baseline oxygen consumption rate (OCR) was measured. Sequential injections of inhibitors (final concentration 4mM ADP, 2.5 ug/mL oligomycin, 4uM carbonyl cyanide-4-(trifluoromethoxy)phenylhydrazone, and 4uM antimycin A) were added into each well and changes in OCR were measured. For Complex I-supported respiration, 10mM pyruvate and 2mM malate were given to mitochondria seeded at 2ug/well.

#### Molecular docking calculation

Crystal structures of tetrameric of human SCOT1 (PDB: 3DLX) were subjected to rigid body molecular docking using online platform, SymmDock (Schneidman-Duhovny *et al*., 2005). Without applying distance constraints, the ten highest scoring solutions were analyzed with the biomolecular visualization tool, Pymol. Residues and sidechains surrounding K296 (human 284-286 and 297-299) are shown as yellow spheres because residues 296-296 are unresolved in all solved structures. The putative interface between SCOT1 structures consistently puts regions surrounding K296 of opposite tetramers nearby (within 42Å distance in 6 of the top 10 solutions).

#### Cytochrome C Oxidase and Citrate Synthase activity from heart tissue homogenate

Citrate synthase (CS) and Cytochrome C Oxidase (COX) activities were determined in heart tissue homogenates at 30°C using a protocol adapted from previously published methods (Spinazzi et al., 2012). Briefly, for CS activity frozen cardiac tissue (25mg) was homogenized at 4°C in CelLytic™ MT Cell Lysis Reagent (C3228) and then incubated on ice for 30 minutes. The samples were then centrifuged at 10,000 x g for 10 minutes at 4°C and protein concentration determined. CS activity was analyzed in 100mM Tris buffer containing 0.1% Triton X-100, pH 8.0 as previously described (Spinazzi *et al*., 2012).

For COX activity, 25mg of cardiac tissue was homogenized at 4°C in 50 mM potassium phosphate buffer containing 1 mM EDTA, and 0.1%Triton X-100 (pH 7.4; final concentration, 5 mg tissue/mL). The samples were incubated on ice for 30 minutes followed by centrifugation at 10,000 x g for 10 minutes at 4°C. COX activity was analyzed in 100mM potassium phosphate buffer as previously described (Spinazzi *et al*., 2012).

### QUANTIFICATION AND STATISTICAL ANALYSIS

#### Assessment of cardiac and mitochondrial function

The numbers of independent experiments are specified in the relevant figure legends. Data are expressed as the mean ± standard error of the mean (SEM). Statistical comparisons between two groups were conducted by unpaired, two-tailed Student’s *t*-test. P<0.05 was considered to be statistically significant. Statistical analysis was performed with Prism 9.0 software (GraphPad).

#### LC-MS/MS data analysis

The LC-MS/MS results were analyzed for identification and quantitation of iqPIR cross-linked peptide pairs as previously described (Chavez *et al*., 2020a). Briefly, LC-MS/MS data raw files were converted to .mzXML format using the ReADW tool in the Trans Proteomic Pipeline software suite (Keller et al., 2005). Mango (Mohr et al., 2018) was used to search for PIR mass relationships and Comet (Chavez et al., 2013) was used to search the mzXML files against the mouse Mitocarta 2 database (Calvo et al., 2015) consisted of both forward and reverse protein sequences (2,084 total sequences). The resulting pepXML files were then analyzed with XLinkProphet (Keller et al., 2018) and filtered for estimated 1% FDR at the non-redundant cross-link level. For dead-end peptides, PeptideProphet (Keller et al., 2002) was applied to filter Comet search results at < 1% FDR. All cross-links passing the threshold were used for quantitation. Light (SH) and heavy (RH) isotope peptide precursors and their fragment peaks containing the stump group were deconvoluted and their peak intensities were extracted to calculate Log2 ratios. The final ratio for each cross-linked peptide pair is the normalized mean value from all contributing ion ratios to the same cross-link from different charge states, different scans, and separate replicate runs with outliers removed, where normalization is achieved by subtracting from each cross-link Log2 ratio the median value of all cross-link Log2 ratios. Finally, the cross-link Log2 ratio p-value reflecting its likelihood of being 0 is calculated based on a statistical t-test using the Log2 ratio mean value, standard deviation, and number of contributing quantified ions. Protein quantitation was estimated by combining together normalized Log2 ratios of all intra-protein cross-links and dead-end peptides corresponding to the protein. The mean Log2 ratio of all contributing ratios (intra-protein cross-link and dead-end) was used after excluding outliers.

## Notes

http://xlinkdb.gs.washington.edu/xlinkdb/index.php

### Competing Interest Statement

The authors have declared no competing interest.

